# Therapeutic Targeting of Measles Virus Polymerase with ERDRP-0519 Suppresses All RNA Synthesis Activity

**DOI:** 10.1101/2020.09.23.311043

**Authors:** Robert M. Cox, Julien Sourimant, Mugunthan Govindarajan, Michael G. Natchus, Richard K. Plemper

**Affiliations:** Institute for Biomedical Sciences, Georgia State University, Atlanta, GA; Emory Institute for Drug Development, Emory University, Atlanta, GA

**Author notes:** these authors contributed equally to this study (Robert M. Cox, Julien Sourimant).

## Abstract

Morbilliviruses, such as measles virus (MeV) and canine distemper virus (CDV), are highly infectious members of the paramyxovirus family. MeV is responsible for major morbidity and mortality in non-vaccinated populations. ERDRP-0519, a pan-morbillivirus small molecule inhibitor for the treatment of measles, targets the morbillivirus RNA-dependent RNA-polymerase (RdRP) complex and displayed unparalleled oral efficacy against lethal infection of ferrets with CDV, an established surrogate model for human measles. Resistance profiling identified the L subunit of the RdRP, which harbors all enzymatic activity of the polymerase complex, as the molecular target of inhibition. Here, we examined binding characteristics, physical docking site, and the molecular mechanism of action of ERDRP-0519 through label-free biolayer interferometry, photoaffinity cross-linking, and *in vitro* RdRP assays using purified MeV RdRP complexes and synthetic templates. Results demonstrate that unlike all other mononegavirus small molecule inhibitors identified to date, ERDRP-0519 inhibits all phosphodiester bond formation in both *de novo* initiation of RNA synthesis at the promoter and RNA elongation by a committed polymerase complex. Photocrosslinking and resistance profiling-informed ligand docking revealed that this unprecedented mechanism of action of ERDRP-0519 is due to simultaneous engagement of the L protein polyribonucleotidyl transferase (PRNTase)-like domain and the flexible intrusion loop by the compound, pharmacologically locking the polymerase in pre-initiation conformation. This study informs selection of ERDRP-0519 as clinical candidate for measles therapy and identifies a previously unrecognized druggable site in mononegavirus L polymerase proteins that can silence all synthesis of viral RNA.

**Importance:** The mononegavirus order contains major established and recently emerged human pathogens. Despite the threat to human health, antiviral therapeutics directed against this order remain understudied. The mononegavirus polymerase complex represents a promising drug target due to its central importance for both virus replication and viral mitigation of the innate host antiviral response. In this study, we have mechanistically characterized a clinical candidate small-molecule MeV polymerase inhibitor. The compound blocked all phosphodiester bond formation activity, a unique mechanism of action unlike all other known mononegavirus polymerase inhibitors. Photocrosslinking-based target site mapping demonstrated that this class-defining prototype inhibitor stabilizes a pre-initiation conformation of the viral polymerase complex that sterically cannot accommodate template RNA. Function-equivalent druggable sites exist in all mononegavirus polymerases. In addition to its direct anti-MeV impact, the insight gained in this study can therefore serve as a blueprint for indication spectrum expansion through structure-informed scaffold engineering or targeted drug discovery.

## Introduction

Morbilliviruses belong to the paramyxovirus family of highly contagious respiratory RNA viruses with negative polarity genomes. The archetype of the morbillivirus genus, MeV, is the most infectious pathogen identified to date with primary reproduction rates of 12-18 (1, 2). Although measles is a vaccine-preventable disease, MeV remains responsible for approximately 100,000 deaths annually worldwide and endemic transmission persists in large geographical regions. Due to its exceptional contagiousness, MeV is typically the first pathogen to reappear when vaccination coverage drops in an area (3, 4). In the aftermath of parental concerns about vaccination safety, four European countries, Albania, Czechia, Greece and the United Kingdom, have regressed to pre-measles eradication status (5). A massive surge in global measles cases is feared as a result of the SARS-CoV-2 pandemic, since immunization campaigns have been suspended as part of the COVID-19 response (6-8). Intensifying preparedness to mitigate the mounting problem is imperative. Effective anti-MeV therapeutics may aid by providing avenues to improved disease management and rapid outbreak control (9).

Towards identifying applicable measles therapeutics, we have developed a MeV inhibitor from high-throughput screening hit to orally bioavailable clinical candidate (10-13). The optimized lead compound, ERDRP-0519, showed pan-morbillivirus inhibitory activity (10), which opened an opportunity for informative efficacy testing in a surrogate assay for human morbillivirus disease. Whereas MeV is human-tropic without natural animal reservoir, infection of ferrets by canine distemper virus (CDV), a morbillivirus closely related to MeV, recapitulates the hallmarks of human measles in a natural animal host of CDV (14). In contrast to the human situation, however, CDV is invariably lethal in ferrets within 12 to 14 days of infection, providing a definitive efficacy endpoint (14). When oral dosing of ferrets with ERDRP-0519 was initiated therapeutically at the first day of viremia, however, outcome of CDV infection was reversed: all treated animals survived, clinical signs were alleviated, and recoverees mounted a robust adaptive immune response protecting against subsequent re-infection (10).

Mechanistically, ERDRP-0519 blocks morbillivirus polymerase activity (10, 11). Conserved among all mononegaviruses, paramyxoviruses encode RNA-dependent RNA polymerase complexes that consist of a large (L) polymerase protein harboring all enzymatic activities and a mandatory chaperone, the viral phosphoprotein (P) (15-19). In addition to phosphodiester bond formation, L mediates capping of viral mRNAs through polyribonucleotidyltransferase (PRNTase) and methyltransferase (MTase) activities (16, 20). Although P does not contain enzymatic activity of its own, it is an essential cofactor required for proper L folding and establishing physical contact between the P-L polymerases and the viral genome, which consists of a ribonucleoprotein (RNP) complex of genomic RNA encapsidated by the viral nucleocapsid (N) protein (21, 22). Resistance profiling of ERDRP-0519 and earlier developmental analogs of the chemotype against MeV and CDV has identified specific escape hotspots in the RdRP and capping domains of the L protein (10, 12). Although some ERDRP-0519 resistance mutations map to spatially distinct regions of the viral polymerase, a prominent cluster of mutations is located in a highly sequence-conserved area in immediate proximity before and after the catalytic site for phosphodiester bond formation (10, 12), a 4-amino acid GDNQ motif (23). Consistent with the notion that sequence conservation arises from selective pressure on the domain, ERDRP-0519 resistance mutations were associated with strong viral fitness penalty in cell culture and *in vivo* (10).

The ERDRP-0519 resistance fingerprint is unique among all small-molecule mononegavirus polymerase inhibitors characterized to date, including small molecule inhibitors of closely related respiratory syncytial virus (RSV) such as AZ-27 (24-26) and AVG-233 (27), and a structurally distinct inhibitor of morbillivirus and respirovirus polymerases, GHP-88309, that we have recently described (28). These features identified the ERDRP-0519 chemotype as a mechanistically first-in-class compound that may have illuminated a conserved, yet currently uncharted, druggable site in mononegavirus polymerases. Despite this high clinical promise, the molecular mechanism of polymerase inhibition by ERDRP-0519 and its physical target docking pose remained unknown, creating a knowledge gap hampering development of the inhibitor class against other mononegavirus targets. Several structural reconstructions of mononegavirus polymerase have been reported recently that provided essential insight into the organization of rhabdovirus (vesicular stomatitis virus and rabies virus (29, 30)), pneumovirus (RSV (31)), and paramyxovirus (parainfluenzavirus 5 (PIV-5) (32)) polymerase complexes. This information established a framework for structural and mechanistic appreciation of polymerase inhibition by ERDRP-0519.

In this study, we have applied a combined biochemical and proteomics characterization strategy to the problem. This approach has revealed the mechanism of polymerase inhibition through *in vitro* RdRP assays using purified recombinant polymerase complexes and synthetic RNA templates, characterized ERDRP-0519 target affinity and the basis for viral escape through biolayer interferometry, and employed photo-affinity labeling for proteomics-based mapping of the physical target site. Through correlation of these independent datasets, we have developed a docking pose for ERDRP-0519 that defines a novel mechanistic paradigm of mononegavirus polymerase inhibition. This information allows pharmacophore-guided tuning of the ERDRP-0519 chemotype if called-for by final de-risking prior to clinical development and outlines a blueprint for the structure-informed identification of novel drug candidates directed against mononegavirus polymerases outside of the ERDRP-0519 indication spectrum.

## Results

Several cryo-electron microscopy-based reconstructions of mononegavirus polymerase proteins have been reported, including structural models of PIV-5 and RSV L polymerases that are closely related to MeV L (31, 32). We have previously identified signature resistance hot-spots for ERDRP-0519 in MeV and CDV L through viral adaptation (10, 12). Escape sites locate to the polymerase and capping domains of L, but notably do not overlap with the resistance profile of recently developed GHP-88309 (28), the only other well-characterized small molecule inhibitor of MeV L (figure 1A).

**Figure 1.**
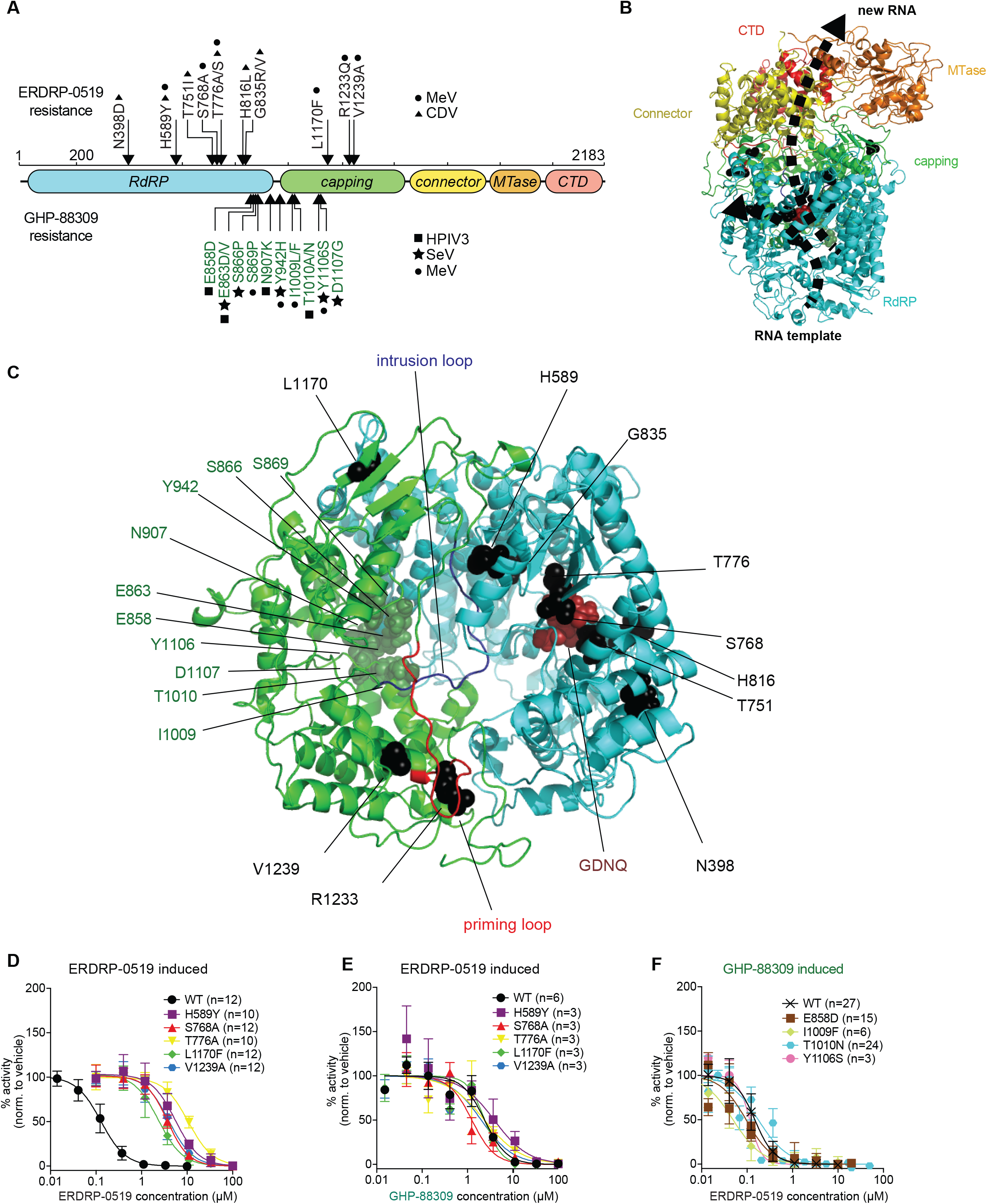
Resistance profiling of ERDRP-0519 and GHP-88309 against MeV polymerase. **A)** 2D-schematic of the MeV L protein. ERDRP-0519 (black) and GHP-88309 (green) resistance mutations are shown. Symbols denote the virus in which each resistance mutation was found (circle, MeV; triangle, CDV; square, HPIV-3; star, Sendai virus). **B)** 3D-homology model of the MeV L polymerase based on the structure of PIV-5 (PDBID: 6v85). The RdRP (cyan), capping (green), connector (yellow), methyltransferase (MTase; orange), and C-terminal (CTD; red) domains are shown. The GDNQ active site is highlighted by red spheres. RNA channels are shown as dotted black lines. **C)** Spatial organization of GHP-88309 and ERDRP-0519 resistance mutations. Shown are locations of tightly clustered GHP-88309 resistance mutations (green) and ERDRP-0519 resistance mutations (black) in the MeV L RdRP (cyan) and capping (green) domains. GDNQ is shown as dark red spheres. Predicted “priming” and “intrusion” loops (32) are shown. **D-F)** Characterization of resistance mutations in a cell-based MeV minigenome assay. Assessed were ERDRP-0519 induced mutations against ERDRP-0519 (D) and GHP-88309 (E), and GHP-88309 induced mutations against ERDRP-0519 (F). Results for the mutants in (F) against GHP-88309 are summarized in (28). Symbols show sample means, curves represent 4-parameter variable slope regression models, EC_50_ values are presented in Table 1. The number of biological repeats (n) is specified for each construct

### Pharmacophore of ERDRP-0519 class is unique

To place resistance mutations in a structural context, we generated a homology model for MeV L based on the PIV-5 L coordinates (figure 1B) and highlighted escape sites of ERDRP-0519 and GHP-88309 (figure 1C). For further comparison, we also located the MeV L homologs of the known resistance sites to RSV L inhibitors AZ-27, RSV L residue 1631 (26), and an RSV L capping inhibitor series, RSV L residues 1269, 1381, and 1421 (33) (supplementary figure 1). Sites associated with resistance to ERDRP-0519 broadly line the internal walls of the central L cavity that houses the catalytic site for polymerization and in which template, substrate, and product exit channels converge. AZ-27 and capping inhibitor resistance sites are located distal from the core cavity, forming part of the flexible interface that links the capping and C-terminal methyltransferase domains. In contrast, GHP-88309 resistance hot-spots are likewise positioned in the core RdRP domain, but these sites were predicted to cluster in the template channel itself rather than the central cavity. ERDRP-0519 and GHP-88309 resistance maps therefore suggested non-overlapping docking poses with MeV L.

To test this hypothesis, we rebuilt ERDRP-0519 and GHP-88309 resistance mutations individually in an MeV minigenome system and examined cross-resistance in dose-response assays. ERDRP-0519-induced mutations reduced MeV polymerase sensitivity to the drug significantly (table 1), independent of whether they had originally emerged from MeV or CDV adaptation (figure 1D). However, none of the ERDRP-0519 resistance mutations altered polymerase susceptibility to GHP-88309 (figure 1E), and none of the previously confirmed substitutions mediating escape from GHP-88309 (28) affected MeV L inhibition by ERDRP-0519 (figure 1F). The lack of cross-resistance between ERDRP-0519 and GHP-88309 confirms that pharmacophores of these inhibitor classes are distinct.

**Table 1.** EC_50_ concentrations against MeV L of resistance mutations induced by viral adaptation to ERDRP-0519 and GHP-88309, respectively.

| L mutation | ERDRP-0519 |  |  | GHP-88309 |  |  |
| --- | --- | --- | --- | --- | --- | --- |
|  | EC<sub>50</sub> [μM] | 95% confidence intervals [μM] | fold change to WT | EC<sub>50</sub> [μM] | 95% confidence intervals [μM] | fold change to WT |
| WT | 0.13 | 0.11 - 0.15 | - | 2.40 | 1.94 - 2.98 | - |
| H589Y<sup>a</sup> | 5.54 | 4.62 - 6.64 | 44.6 | 3.91 | 1.87 - 8.17 | 1.8 |
| S768A<sup>a</sup> | 3.72 | 3.35 - 4.12 | 29.1 | 1.26 | 0.93 - 1.71 | 0.6 |
| T776A<sup>a</sup> | 10.67 | 9.41 - 12.10 | 76.6 | 2.39 | 1.45 - 3.94 | 1.1 |
| L1170F<sup>a</sup> | 2.36 | 1.99 - 2.79 | 16.9 | 2.41 | 1.70 - 3.4 | 0.7 |
| V1239A<sup>a</sup> | 3.97 | 3.34 - 4.72 | 29.8 | 2.03 | 1.34 - 3.07 | 0.7 |
| E858D<sup>b</sup> | 0.10 | 0.08 - 0.11 | 0.7 | 15.04* | 10.69 - 19.68* | 6.7 |
| I1009F<sup>b</sup> | 0.05 | 0.04 - 0.07 | 0.3 | 18.62* | 13.78 - 25.51* | 8.3 |
| T1010N<sup>b</sup> | 0.18 | 0.15 - 0.21 | 1.5 | 8.36* | 4.40 - 15.71* | 3.8 |
| Y1106S<sup>b</sup> | 0.11 | 0.09 - 0.14 | 0.6 | >100* | >100* | >50 |

\*reported previously (28)
<sup>a</sup> resistance mutation induced by ERDRP-0519
<sup>b</sup> resistance mutation induced by GHP-88309

### Target binding affinity of ERDRP-0519 matches inhibitory concentration

For biochemical positive target identification, we expressed and purified recombinant standard and mutant MeV P-L complexes from insect cells (figure 2A). Following *in vitro* mono-biotinylation of the protein preparations, complexes were immobilized on streptavidin high-density biolayer interferometry (BLI) probes and ERDRP-0519 association and dissociation curves recorded at increasing compound concentrations (figure 2B-E). Analysis of binding kinetics to standard MeV P-L revealed dissociation constants (K_D_) of 140 nM, which resembles the EC_50_ of 230 nM that we had recorded for ERDRP-0519 in cell-based anti-MeV assays (10).

**Figure 2.**
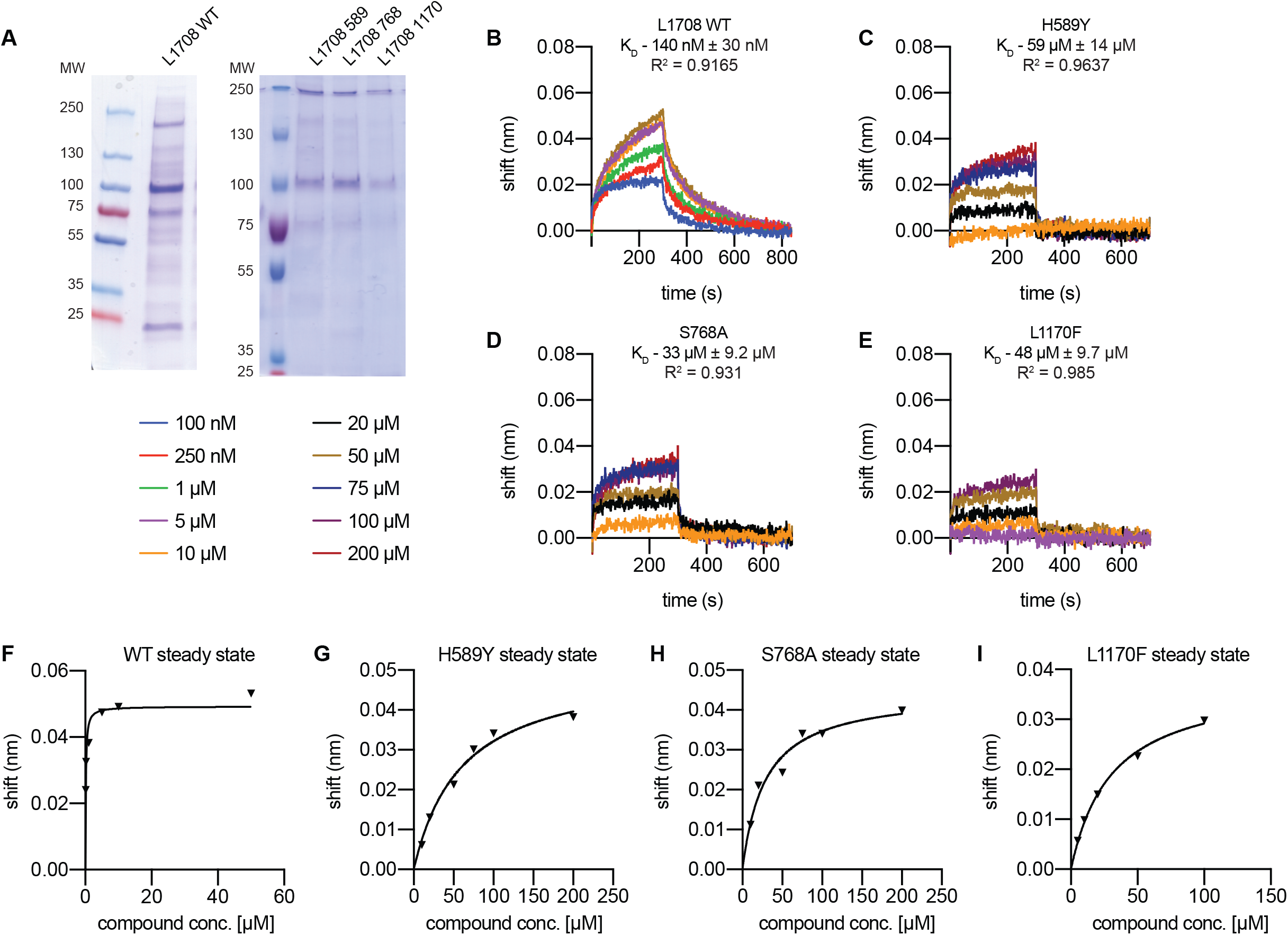
Target binding affinity of ERDRP-0519. **A)** SDS-PAGE of purified MeV L_1708_ (28) used for BLI studies. **B-E)** BLI of ERDRP-0519 and purified standard (WT) MeV L and L harboring selected ERDRP-0519 resistance mutations. K_D_ values and goodness of fit are shown for each construct. **F-I)**, Steady-state analyses of BLI binding saturation in (B-E). Concentration-dependent steady-state BLI sensor response signals were plotted for the different L populations (WT or carrying resistance mutations).

To test whether resistance mutations impair ligand interaction with the L target, we subjected mutant P-L complexes harboring a substitution in one of three distinct resistance clusters in the linear L sequence, H589Y, S768A or L1170F, respectively, to BLI analysis. These mutations increased KD values to 59, 33, and 48 µM, respectively, which resembles an approximate 35- to 63-fold drop in target affinity (figure 2C-E). In addition, ERDRP-0519 binding to standard MeV L reached saturation at low micromolar concentrations (figure 2F), whereas binding to each of the mutant L proteins was drastically reduced and saturation was not reached at the highest concentration tested, 200 μM (figure 2G-I). These BLI results positively identify the MeV L protein as ERDRP-0519 target and demonstrate that viral resistance originates from reduced ligand binding affinity.

### ERDRP-0519 resistance mutations restore initiation of RNA synthesis but not RNA elongation

For mechanism-of-action characterization, we applied purified recombinant MeV P-L complexes to *in vitro* RdRP assays in the presence and absence of ERDRP-0519, using _32_P tracers and either a synthetic 16-mer RNA template containing an MeV-specific promoter region to monitor *de novo* initiation of RNA synthesis at the promoter (figure 3A) or a synthetic primer-template pair to assess RNA elongation by a committed polymerase (figure 3B). Autoradiograms and phosphoimager-based quantitations revealed efficient and dose-dependent inhibition of standard MeV polymerase complexes under either assay condition with IC_50_ values of 0.15 µM and 0.1 µM, respectively (figure 3C-F). These active concentrations closely recapitulated the EC_50_ of ERDRP-0519 against MeV in cell-based assays. A catalytically defective L mutant harboring an N774A substitution in the polymerase active site confirmed that RNA synthesis was MeV P-L specific and not due to co-purified cellular contaminants.

**Figure 3.**
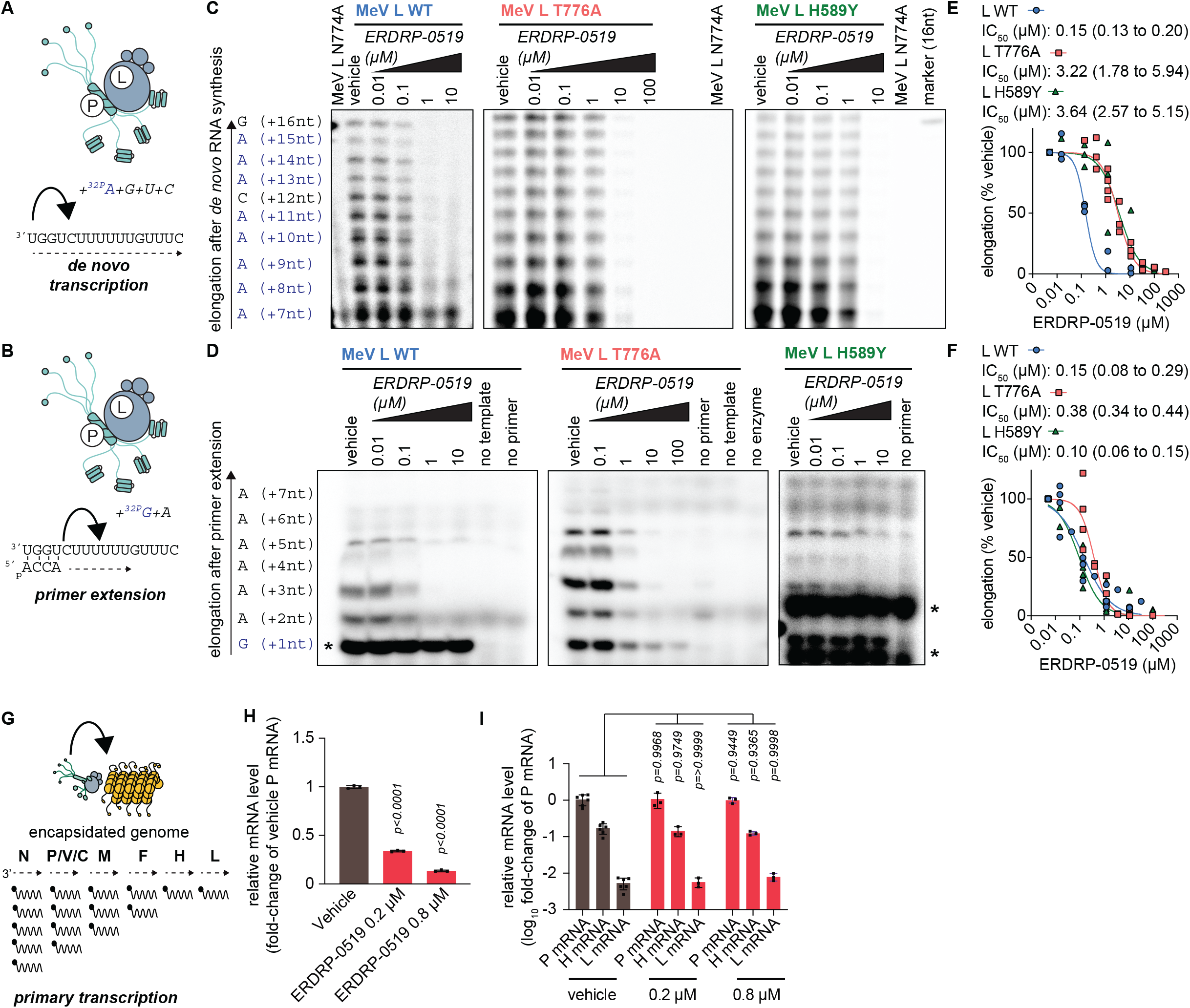
ERDRP-0519 potently inhibits *de novo* RNA synthesis. **A-F)** Purified recombinant WT MeV L-P complexes or complexes harboring the L_T776A_ or L_H589Y_ resistance mutations were incubated with either a 16-nt RNA template (A) and the represented NTPs to assess *de novo* RNA synthesis or a 16-nt RNA template (B), a 5’-phosphorylated 4-nt primer, and the represented NTPs to assess primer extension. Representative autoradiograms are shown for *de novo* initiation (C) and primer extension (D). Purified L-P complexes with an L_N774A_ substitution in the catalytic GDNQ motif (59) served as control for specificity of the *de novo* initiation assay. Primer extension assays were performed in the absence of primer, template, or enzyme to control for contaminating *de novo* RNA synthesis driven directly by the template. Densitometry analysis was performed on elongation products of 15 to 16-nt in length (E) or 7 to 9-nt in lengths (F) and represent n = 3-4 biological repeats. EC_50_ values represent 4-parameter variable slope regression models, 95% confidence intervals are shown. Positions of unspecific background signals are indicated with *. **G-I)** Effect of ERDRP-0519 incubation on MeV primary transcription, represented in (G). Cells were infected with recMeV-Anc (MOI = 3) and incubated with vehicle (0.1% DMSO) volume control, or 0.2 µM or 0.8 µM ERDRP-0519. Infected cells were harvested 4 hours after infection. Incubation with ERDRP-0519 significantly decreased relative P-encoding mRNA amounts 4 hours post infection (H), but did not alter steepness of the MeV primary mRNA transcription gradient (I). Symbols represent individual biological repeats (n = 3), graphs show sample means. Statistical analysis through two-way ANOVA with Dunnett’s multiple comparison post-hoc test, P values are specified.

Polymerase complexes containing an ERDRP-0519 resistance mutations H589Y or T776A showed greatly reduced susceptibility to compound-mediated suppression of *de novo* polymerase initiation, reflected by inhibitory concentrations of 3.64 µM and 3.22 µM, respectively, which represents a 24- and 21-fold increase in IC_50_ value (figure 3C,E). By comparison, the same resistance mutations had a marginal effect on lifting ERDRP-0519 inhibition of RNA elongation in the *in vitro* RdRP assay. The H589Y substitution did not increase the IC_50_ concentration at all (IC_50_ 0.10 µM) and T776A caused only a moderate 2.5-fold IC_50_ increase (IC_50_ 0.38 µM) (figure 3D,F). Because these resistance mutations mediate robust viral escape in cell-based inhibition assays, the *in vitro* RdRP results highlight inhibition of RNA synthesis at the promoter as the physiologically relevant antiviral effect of ERDRP-0519. Resistance mutation-insensitive inhibition of primer extension is very likely a consequence of the artificial conditions of the *in vitro* assay using a non-encapsidated template that does not fully recapitulate the natural system.

To test this hypothesis, we assessed the effect of ERDRP-0519 on the relative amount of viral mRNA transcripts synthesized in infected cells during the first 4 hours after infection, representing the primary transcription phase (figure 3G). MeV P mRNA levels were dose-dependently reduced by ERDRP-0519 (figure 3H). Importantly, however, the inhibitor had no significant effect on the ratio of different viral mRNAs relative to each other (figure 3I), the viral transcription gradient. This finding indicates that ERDRP-0519 is unable to inhibit RNA elongation by an engaged polymerase complex in infected cells, which would steepen the transcription gradient.

### ERDRP-0519 blocks all phosphodiester bond formation

All allosteric paramyxovirus and pneumovirus polymerase inhibitors subjected to *in vitro* RdRP assays to date were found unable to block the formation of the first phosphodiester bond during backpriming and/or primer extension (27, 28, 34). Backpriming refers to the spontaneous formation of a circular hairpin structure of the non-encapsidated synthetic templates (35) that allows extension of the resulting paired 3’-ends beyond the actual length of the template (figure 4A). To assess whether the same limitation applies to ERDRP-0519, we applied the MeV polymerase complex to a previously described 25-mer RNA template derived from an authentic RSV promoter sequence that is capable of efficient backpriming (35). The RSV template was efficiently recognized as a suitable substrate for *de novo* RNA synthesis by the MeV polymerase (figure 4B). Potency of inhibition of *de novo* initiation by ERDRP-0519 resembled that observed for the MeV promoter (IC_50_ 0.12 vs 0.15 µM). In contrast to all available inhibitors formerly described, however, all 3’-elongation after backpriming was equivalently dose-dependently blocked (figure 4C).

**Figure 4.**
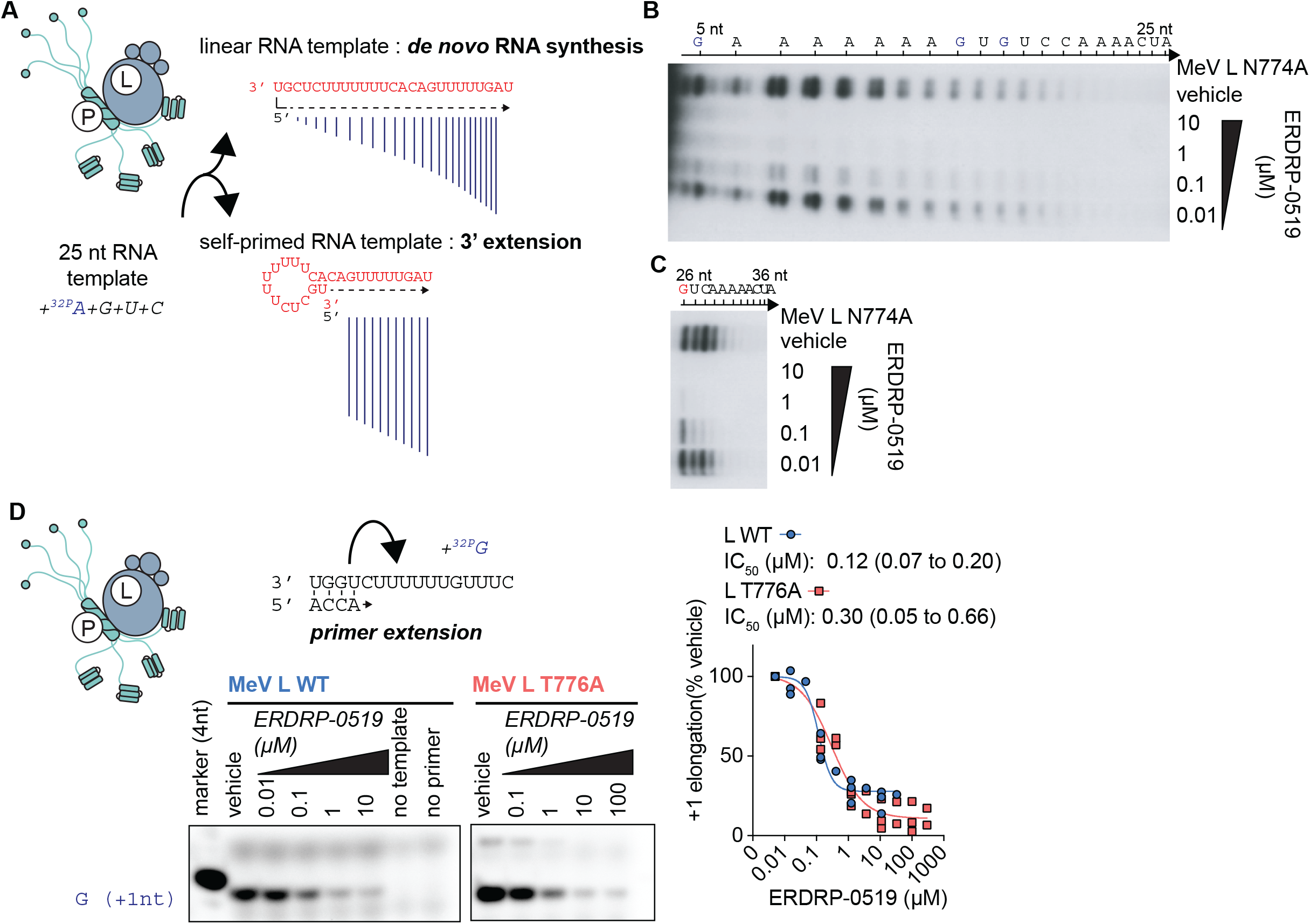
Effects of ERDRP-0519 on single nucleotide addition. **A-C)** Purified recombinant P-L complexes were incubated with the specified NTPs and a 25-nt RNA template driving both *de novo* initiation and back-priming (35) (A). A representative autoradiogram (n = 3) is shown, divided for clarity into dose-dependent inhibition of *de novo* RNA synthesis initiation at the promoter by ERDRP-0519 (B) and inhibition of 3’-elongation after back-priming (C). Products of less than 5-nt are not perceptible due to background from unincorporated ^32^P-labelled nucleotides. **D)** Dose-dependent inhibition of primer extension by ERDRP-0519 as in figure 3B, but only GTP was added to visualize incorporation of the first nucleotide. EC_50_ values represent 4-parameter variable slope regression models, 95% confidence intervals are shown (n = 3).

For validation of this unprecedented finding, we carried out a primer extension assay in the absence of all NTPs but the ^32^P-GTP tracer, consequently allowing monitoring incorporation of the first nucleotide, specifically (figure 4D). Again, ERDRP-0519 dose-dependently blocked incorporation of the very first nucleotide with equivalent potency for standard (L_WT_; IC_50_ 0.12 vs 0.15 µM) and resistant (L_T776A_; IC_50_ 0.3 vs 0.39 µM) MeV polymerases to inhibition of multi-nucleotide elongation (figure 4B).

In conclusion, these results consistently demonstrate that ERDRP-0519 directly suppresses all phosphodiester bond formation, setting the compound mechanistically apart from all other allosteric mononegavirus inhibitor classes characterized to date.

### Photoaffinity labeling maps the ERDRP-0519 target site to the central L cavity

To map the physical target site of ERDRP-0519, we synthesized a photo-activatable analog of the compound, ERDRP-0519_az_, through installation of an aryl azide moiety at C-2 position of the ERDRP-0519 piperidine ring via a short tether (figure 5A). Analog design was guided by our extensive insight into the structure-activity relationship (SAR) of the ERDRP-0519 chemotype that we had acquired in previous work (11). Bioactivity testing of ERDRP-0519_az_ against MeV revealed dose-dependent suppression of virus replication and an EC_50_ of 12.1 µM (figure 5B). No cytotoxicity was detectable at the highest concentration tested (100 µM), indicating specific virus inhibition by the photo-activatable probe. Compared to ERDRP-0519, however, antiviral potency of ERDRP-0519_az_ was reduced approximately 50-fold, which may reflect reduced plasma membrane permeability of the azide analog in cell-based assays or partially impaired target access of the modified compound. We compensated for this reduction in activity by incubating the P-L complexes in the presence of 40 µM compound prior to photo-activation.

**Figure 5.**
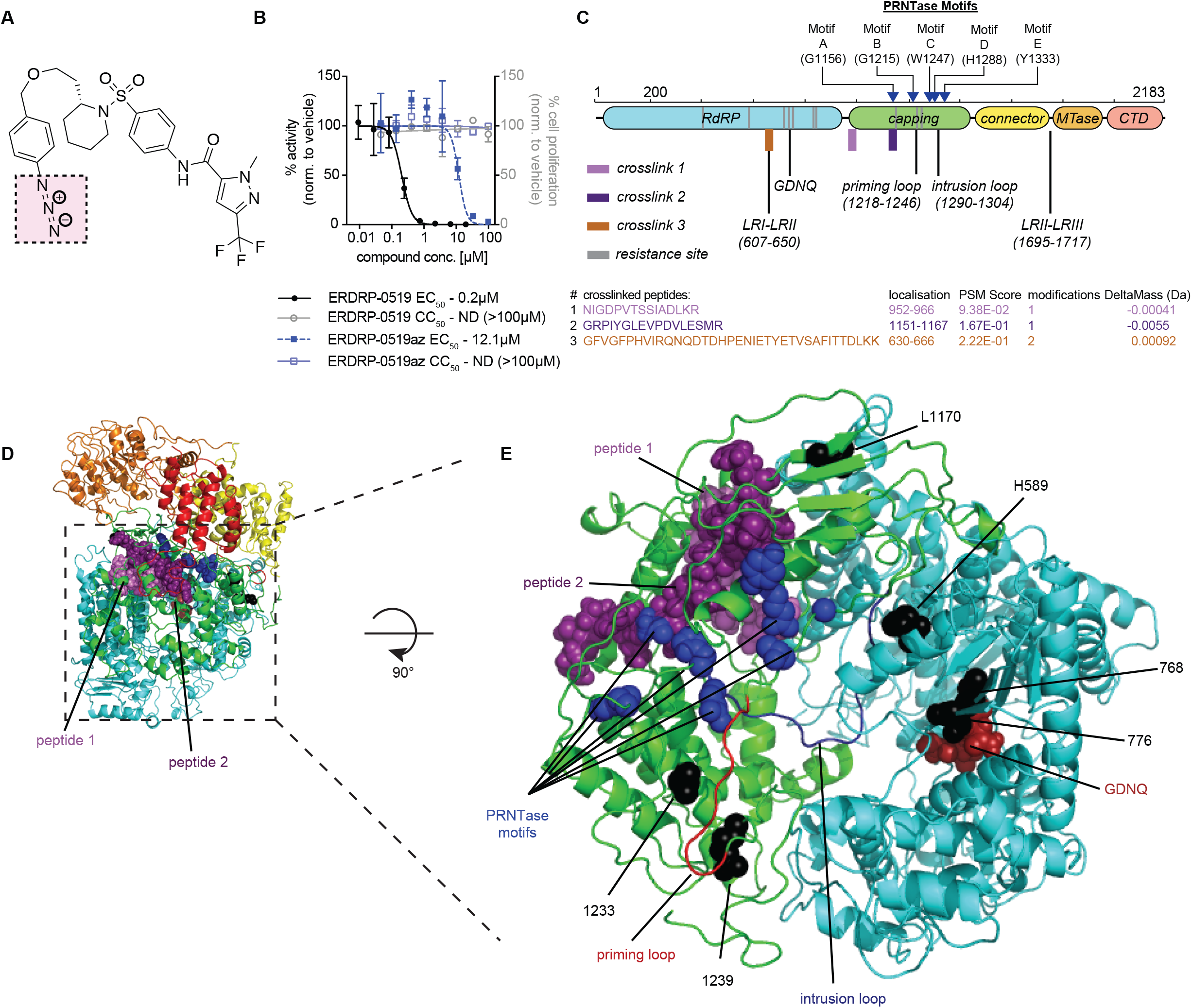
Photoaffinity labeling-based target mapping of ERDRP-0519. **A)** Structure of photoactivatable compound ERDRP-0519_az_; the reactive azide moiety is highlighted (pink square). **B)** ERDRP-0519_az_ is bioactive and displays no appreciable cytotoxicity. Symbols show means of three biological repeats, graphs represent 4-parameter variable slope regression models where possible. EC_50_ and CC_50_ are shown. **C)** 2D-schematic of the MeV L protein with locations of crosslinked peptides identified by photoaffinity labeling (top). The RdRP (cyan), capping (green), connector (yellow), MTase (orange), and C-terminal (light-red) domains, locations of known unstructured regions (LRI-LRII; LRII-LR-III), the GDNQ active site, specific amino acids motif in the capping domain, and positions of the intrusion and priming loops are shown. Sequences of peptides engaged by ERDRP-0519_az_ (bottom). Specified are peptide location, spectrum match (PSM) score, number of ligands present, and delta mass. **D)** Location of peptides 1 (pink) and 2 (purple) in MeV L. **E)** Close-up top view of the capping and RdRP domains showing the adjacent locations of peptides 1 and 2. The priming loop (red), intrusion loop (blue), PRNTase motifs (blue spheres) and GDNQ active site (red spheres) are shown.

Photo-coupling of ERDRP-0519_az_ to purified L followed by LC-MS/MS analysis after trypsin digestion of the ligand-L complexes identified three discrete peptides that are located in the capping and RdRP domains of the L protein (figure 5C). Peptides 1 and 2 (spanning residues 952-966 and 1151-1167, respectively) are located in immediate proximity to each other, forming part of the upper wall of the central polymerase cavity (figure 5D). Peptide 3 spans residues 630-666 in the RdRP domain, a variable and unstructured region in mononegavirus polymerases. However, this peptide is part of a variable region in the viral polymerase (supplementary figure 2) and carried simultaneously two covalently bound ERDRP-0519_az_ moieties, rendering it a very likely candidate for nonspecific crosslinking.

We therefore concentrated further analysis on peptides 1 and 2, which are near the intersection between the polymerase capping, connector and MTase domains, and in close proximity to highly conserved polymerase motifs such as the proposed PRNTase domain (HR moiety of motif D and G1156 of motif A) as well as both the postulated paramyxovirus L priming and “intrusion” loops (32) (figure 5E).

### Docking predicts that ERDRP-0519 locks the polymerase in pre-initiation conformation

Overlaying the photo-crosslinking results and resistance maps in the MeV L structural model revealed a circular arrangement of all potential ERDRP-0519 anchor points along the interior lining of the central polymerase cavity (figure 6A). Docking of ERDRP-0519 into the L structure was guided by the following constraints: positioning of the ligand is compatible with the formation of covalent bonds with residues in photo-crosslinking defined peptides 1 and 2; and the docked compound is in equal proximity to confirmed resistance sites H589, S768, T776, L1170, R1233, and V1239. In addition to ERDRP-0519, two chemical analogs, the original screening hit 16677 (13) and the first-generation lead AS-136 (36), were used as ligands for *in silico* docking (supplementary figure 3). A conserved top-scoring pose for 16677, AS-136, and ERDRP-0519 was returned that placed each analog in a pocket at the intersection of MeV L capping and RdRP domains, between motifs A and D of the predicted PRNTase domain (figure 6A). This pose places the ligand in approximately 26Å distance to peptide 1 and 2.7-10.4Å distance to peptide 2, establishing hydrogen bond interactions between residue Y1155 in peptide 2 and the central pyrazole ring of the ERDRP-519 scaffold (Figure 6B). Additional hydrogen bonds are predicted between H1288 in the polymerase HR motif located on the postulated intrusion loop of paramyxovirus polymerases (32) and the sulfonyl group of the ERDRP-0519 scaffold, and N1285 of the intrusion loop and the piperidine moiety of ERDRP-0519. Equivalent ERDRP-0519 docking poses could not be identified for PIV-5, RSV, and VSV L (supplementary figure 4), which are all insensitive to the compound.

**Figure 6.**
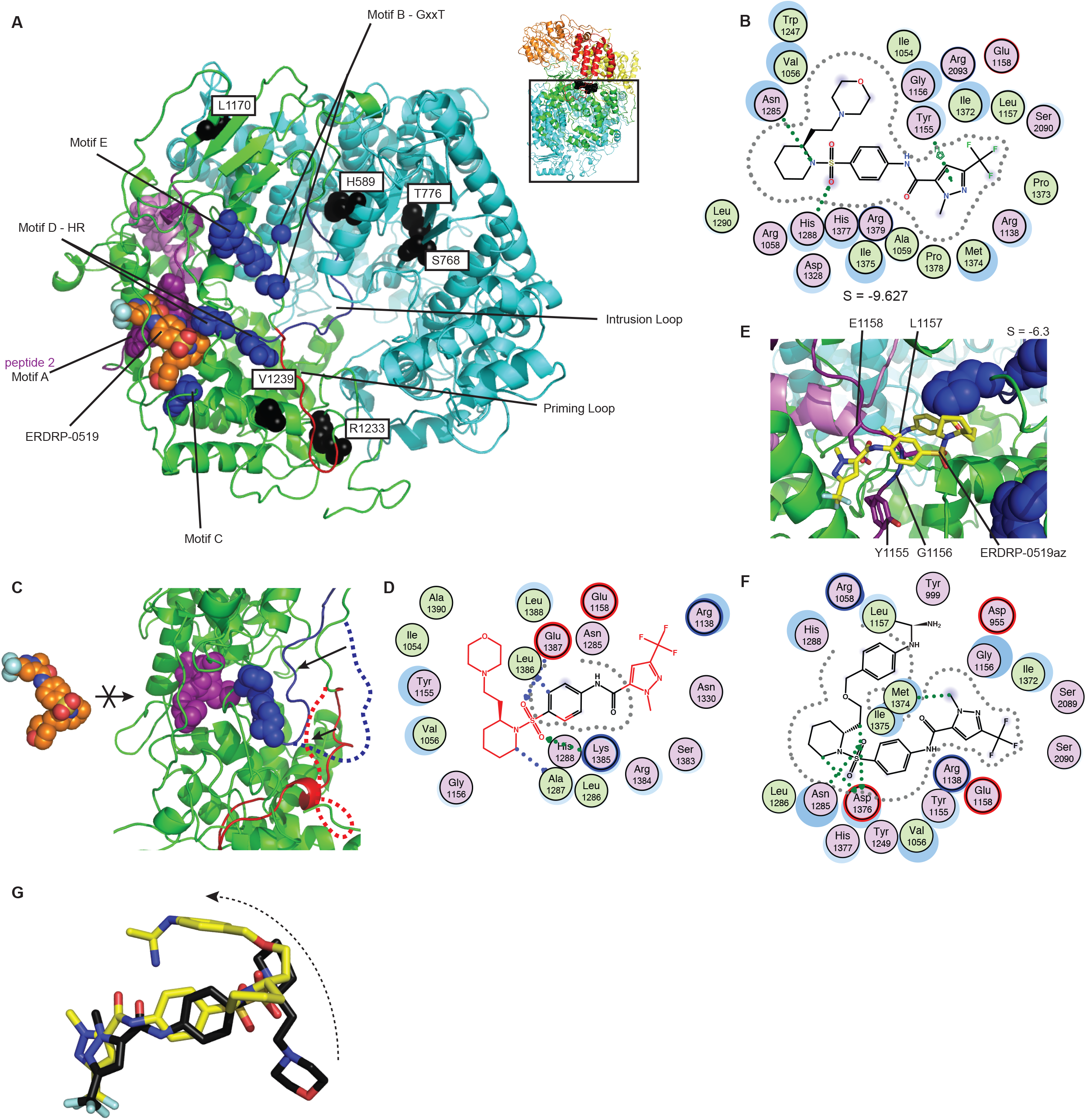
*In silico* docking and ERDRP-0519 pharmacophore extraction. **A)** Top view of the RdRP (cyan) and capping (green) domains of MeV L. PRNTase motifs, ERDRP-0519 resistance mutations, and ERDRP-519 are shown as blue, black, and magenta spheres, respectively. Peptide 2 (purple), the priming loop (red) and intrusion loop (blue) are labeled. The top scoring docking pose places ERDRP-0519 (orange) in close proximity to peptide 2. **B)** 2D-interaction projection of the top scoring ERDRP-0519 docking pose. The sulfonyl oxygen is predicted to interact with H1288 of the HR motif and the pyrazol ring hydrogen bonds with Y1155 of peptide 2. **C-D)** The binding pocket of ERDRP-0519 is not available in an MeV L homology model based on VSV L in initiation conformation (PDBID: 5A22) (C). Attempts to insert ERDRP-0519 result in multiple steric violations (moieties of the compound structure highlighted in red) (D). **E-G)** Close-up view (E), 2D-interaction map (F), and scaffold overlays (G) of the corresponding top-scoring covalent ERDRP-0519_az_ docking pose. Positioning and main scaffold orientation resembles the pose of ERDRP-0519, although the sulfonyl group hydrogen bonds with D1378 instead of H1288. The lateral ring system containing the azide moiety of ERDRP-0519_az_ (yellow sticks) must rotate (G) to fit.

In top-scoring position ERDRP-0519 should lock MeV L intrusion and postulated priming loops in place, blocking reorganization of the central cavity for polymerase initiation. To test this hypothesis, we attempted to dock the ligand into the equivalent space in an MeV L model based on the structure of VSV L (30), in which the priming loop is considered to be in initiation conformation, displacing the intrusion loop from the central polymerase cavity. Only in this configuration can the polymerase accommodate RNA in the cavity (37). However, the reorganization of the PRNTase domain into initiation mode eliminated the ERDRP-0519 binding pocket (figure 6C-D), suggesting that the compound stabilizes a pre-initiation conformation of the polymerase.

To explore whether the docking pose supports covalently binding of photo-reactive ERDRP-0519 to experimentally identified target residues, we examined docking of ERDRP-0519_az_ in the pre-initiation L model, using a covalent *in silico* docking approach. A top-scoring pose resembled that of standard ERDRP-0519 (figure 6E-F). However, the terminal ring system carrying the azide arm was rotated by 180° (figure 6G) and was predicted to covalently engage L1157 in peptide 2. This pose supports peptide 2 as a premier covalent target site for ERDRP-0519_az_. However, ERDRP-0519_az_ did not hydrogen bond with Y1155 and H1288, which may explain the approximately 60-fold decrease in potency of ERDRP-0519_az_ compared to standard ERDRP-0519.

### Ligand-driven 3D-quantitative SAR validates the *in silico* docking-derived pharmacophore

To independently assess the predictive power of the photoaffinity labeling-based docking pose, we selected from our synthetic collection an informative set of 33 analogs of the ERDRP-0519 chemotype with EC_50_ values ranging from 0.005-65 µM (table 2) (11, 36) to generate a ligand-driven 3D-QSAR model. This panel was divided into a training subset of 16 analogs with an EC_50_ range of 0.005 to 23 μM and a 17-member test subset. We next generated an *in silico* library of 25–523 distinct conformations for each compound of the training set and aligned these into 287 initial pharmacophore models that were generated with the AutoGPA module embedded in the MOE software package (38). Individual models were ranked based on goodness of fit (R^2^) and predictive capacity (q^2^) against the training set (figure 7A), leading to the identification of a top-scoring 3D-QSAR model with R^2^ = 0.9749 and q^2^ = 0.7768. Predicted and experimentally determined activity of the training subset were correlated with a regression slope of 0.9803 through the origin. Validation of this model against the 17-analog test subset that included ERDRP-0519, ERDRP-0519_az_, and the first generation lead AS-136 returned an R^2^ of 0.54 and regression slope of 0.89 for the test set of analogs (figure 7A).

**Table 2.**
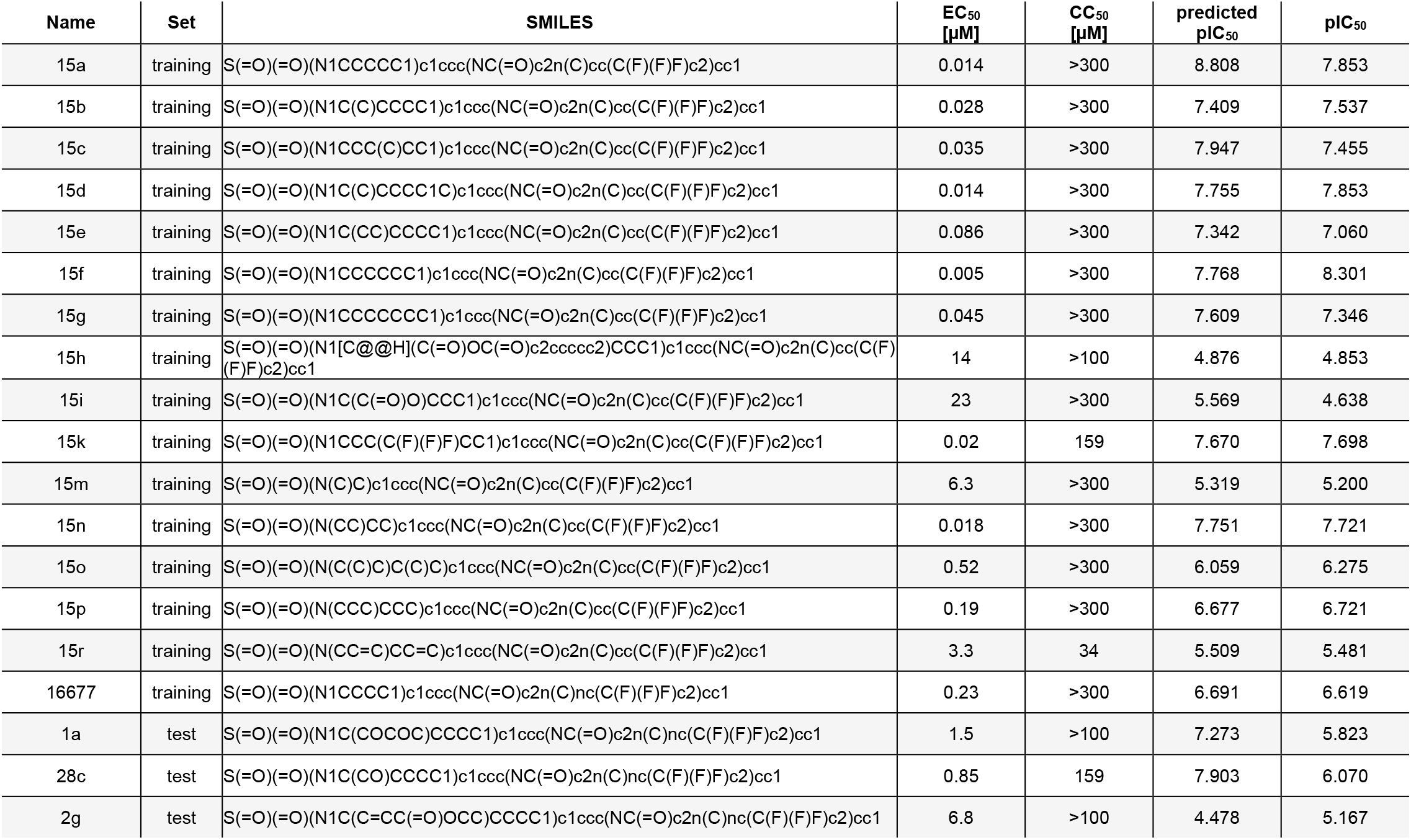
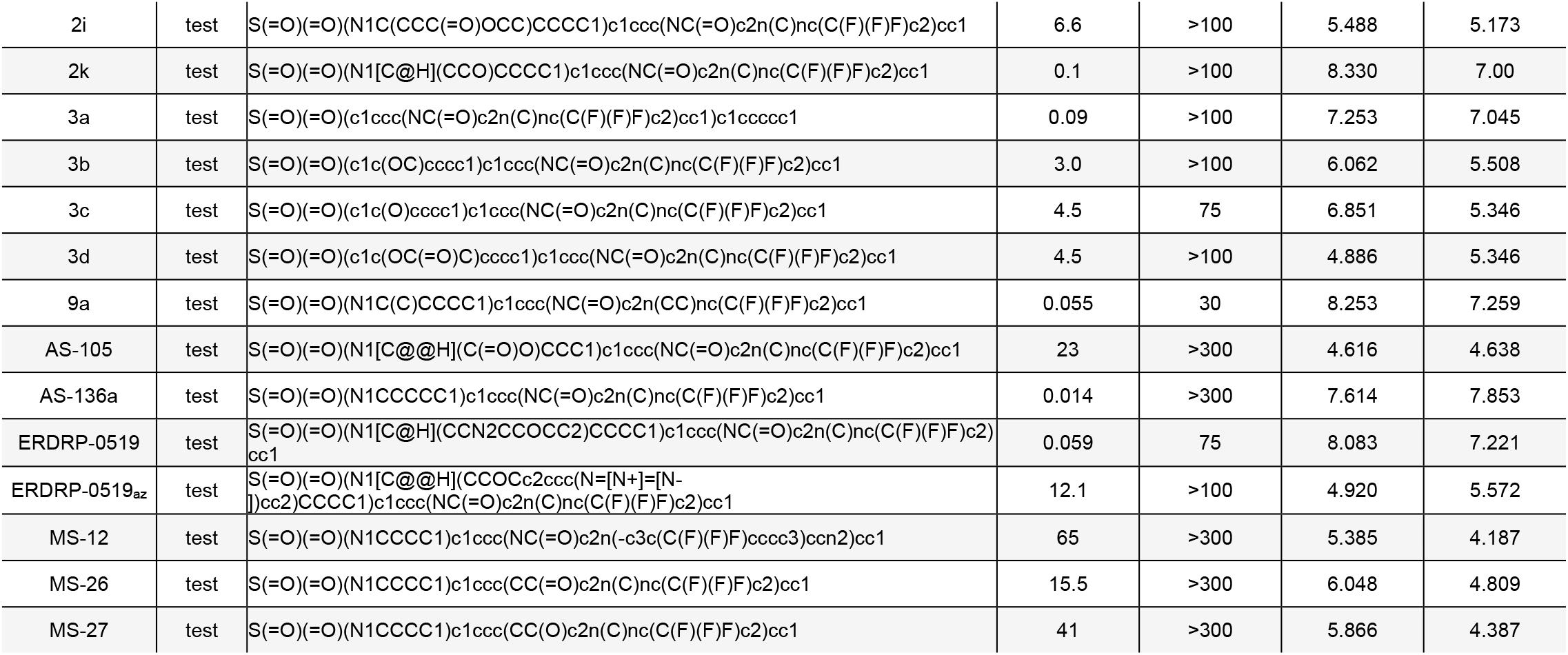
ERDRP-0519 analogs used for development of the 3D-QSAR model. Experimental (11, 36, 58) and predicted activities are shown.

**Figure 7.**
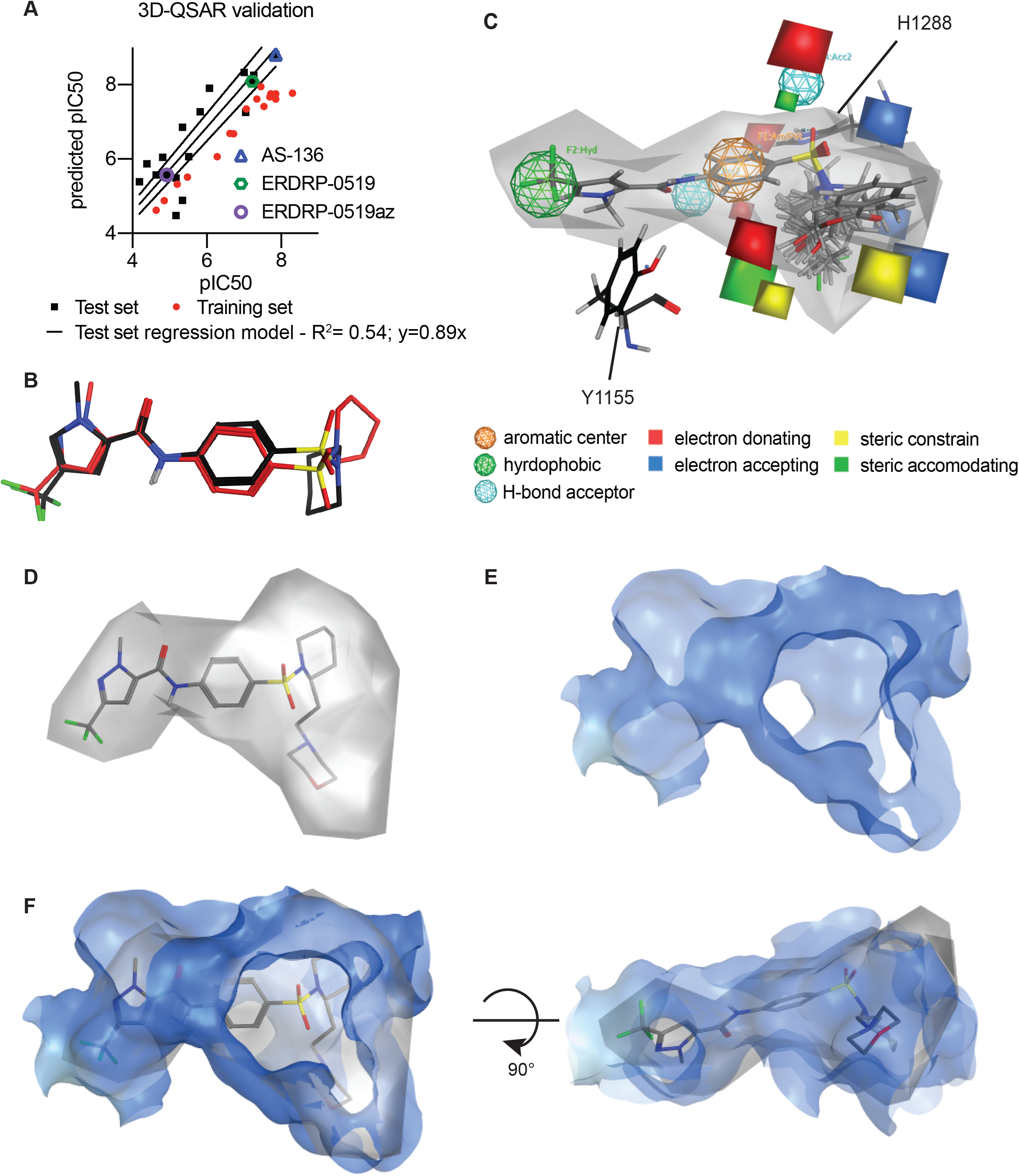
Independent ERDRP-0519 docking validation through ligand-driven 3D-QSAR modeling. **A)** Development of the 3D-QSAR model. Individual data points of the training and test set, goodness of fit (R^2^), and slope of the best fit correlation through the origin are shown for the test set. **B)** Overlays of docking poses of the core ERDRP-0519 scaffold identified by the 3D-QSAR (red) and photocrosslinking-informed *in silico* docking (black). **C)** Graphical representation of the 3D-QSAR model showing space filling (grey) and pharmacophore features of the model. Proximity of features to H1288) and Y1155 is shown. **D-F)** Overlays of the space filling portion of the 3D-QSAR model (grey) with the L-provided binding pocket (blue). Individual views of the space-filling portion (D) and the binding pocket (E), and top and side views of the overlays (F) are shown.

Direct comparison of the 3D-QSAR-derived pharmacophore and the top-scoring docking pose revealed a close overlap of the predicted conformation of the target-bound ERDRP-0519 core scaffold (root-mean-square-deviation (RMSD): 5.484; figure 7B). Most notable of the highly conserved features between both pharmacophores are the predicted strong hydrogen bond interactions between the target pocket and the piperidine and sulfonyl groups of the ERDRP-0519 scaffold (figure 7C). Graphical overlay of both models furthermore revealed a strong correlation between shape and dimension of the available space in the target site illuminated by the L protein structural model and the ligand-occupied space requested by the 3D-QSAR model (figure 7D).

## Discussion

Despite the major clinical threat posed by mononegaviruses and several decades of drug development, ribavirin is the only small-molecule antiviral therapeutic licensed for clinical use against any pathogen of the order to date. However, ribavirin use against RSV has been largely discontinued due to a poly-pharmacological mechanism of action (39-41), pronounced side effects (42, 43), and limited efficacy (44), creating a major unaddressed clinical need. We favor the mononegavirus RdRP complex as a premier druggable target, based on its diverse enzymatic activities, its unique catalytic activity that lacks a cellular equivalent, and its critical importance for both viral replication and the expression of non-structural immunomodulatory viral proteins that counteract the host innate antiviral response (45). Groundbreaking progress in the structural understanding of the organization of mononegavirus polymerase complexes (29-32) has established a foundation to define distinct druggable sites in the L polymerase.

Focusing on the closely related pneumovirus and paramyxovirus families specifically, four distinct non-nucleoside chemotypes have been subjected to defined RdRP assays using purified polymerase complexes and synthetic RNA substrates. Of these, two (AZ-27 (34) and AVG-233 (27)) are RSV L-specific, one (GHP-88309 (28)) inhibits paramyxoviruses of the respiro- and morbillivirus genera, and ERDRP-0519 specifically blocks morbillivirus polymerases (10, 12). Remarkably, all of these compounds have been demonstrated to interfere with *de novo* polymerase initiation at the promoter and backpriming, albeit in the case of AZ-27, AVG-233, and GHP-88309, after incorporation of an additional 2-4 nucleotides (27, 28, 34). This delayed polymerase arrest demonstrates that these compounds do not directly block phosphodiester bond formation. Structural evidence (29-32, 46) and functional characterization (27, 34) rather indicates pharmacological interference with conformational changes of the polymerase as the enzyme transitions from initiation to RNA elongation mode. Consistent with this conclusion, none of these three inhibitor classes affects extension of the RNA template after backpriming (27, 28, 34), which is considered to mimic RNA elongation by a committed polymerase complex (35, 47, 48).

Characterization of ERDRP-0519 demonstrated that preventing the switch to elongation mode is not the only mechanism available to allosteric small molecule inhibitors to block mononegavirus polymerases, since the compound interrupted both initiation at the promoter and RNA elongation after backpriming with equal potency. Underscoring a unique mechanism of action of ERDRP-0519, inhibition of polymerase initiation was furthermore immediate, indicating that all phosphodiester bond formation ceases in the presence of the inhibitor. As a consequence of this MOA principle, ERDRP-0519 potency in the RdRP closely resembled that in cell based antiviral assays. This behavior could not be taken for granted, since other polymerase inhibitor classes showed a remarkable discrepancy between *in vitro* and *in cellula* active concentrations (27, 28, 34). Only target binding affinity of ERDRP-0519 determined through BLI furthermore resembled the inhibitory concentration range, indicating that in the case of ERDRP-0519 interaction with purified P-L complexes assesses compound docking into the primary, bioactivity-relevant target site. By extension, this finding provided support for the physiological relevance of the photoaffinity labeling-based target mapping strategy that likewise relied on *in vitro* interaction of the ligand with purified polymerase complexes.

Although three peptides were identified by photocrosslinking, we primarily focused on peptides 1 and 2. Peptide 3 is located in a highly variable and unstructured region that is completely solvent exposed in the L structural model (supplementary figure 2) and two covalently bound ERDRP-0519_az_ molecules were detected simultaneously on the same peptide 3 molecule, suggesting nonspecific crosslinking events. In contrast, peptides 1 and 2 are separated by some 200 amino acids in the linear polypeptide sequence, but are located in close spatial proximity to each other in a structurally defined section of the native PRNTase domain. Physical engagement of residues in either of these two peptides by ERDRP-0519_az_ in a consistent, specific docking pose was structurally only possible if the compound is positioned in the central cavity of the polymerase complex.

The top-scoring docking pose proximal to peptide 2 offers a mechanistic explanation for how ERDRP-0519 can prevent all phosphodiester bond formation. The lead candidate residues in peptide 2 for covalent interaction with ERDRP-0519_az_, Y1155, G1156, L1157, and E1158, are all located at a critical intersection between the polymerase capping and RdRP domains. Direct interaction between ERDRP-0519 and the PRNTase intrusion and priming loops sterically blocks repositioning of the intrusion loop towards the cavity wall as seen in the VSV, RABV, and RSV L structures (figure 6C; supplementary figure 5) (29-31), thereby obstructing access of template RNA to the cavity and preventing rearrangement of the priming loop into initiation conformation. Pharmacologically locked into a single conformation that cannot accommodate template RNA in the central cavity (32), the polymerase complex is unable to initiate +1 RNA synthesis or elongate RNA in primer extension and after backpriming. In addition, H1288 of the HR moiety in motif D of the PRNTase domain, which mediates the nucleophilic attack on 5′-α-phosphorus atoms in 5′-triphosphorylated RNA to form covalent L-pRNA intermediates (49), is essential for the initiation of L-mediated RNA synthesis (50). Hydrogen bonding between the sulfonyl group of ERDRP-0519 and this residue is very likely catastrophic for the formation of productive initiation complexes.

Confirmed resistance sites to ERDRP-0519 line the internal wall of the central polymerase cavity, but individual hot-spots do not cluster in the native structure and none is predicted to be in direct contact with the docked ligand. Remarkably, however, resistance sites were located in highly sequence conserved L domains such as on the priming and intrusion loops (i.e. H589Y, R1233Q, and V1239A) and/or in immediate proximity of functional motifs (i.e. S768A and T776A framing the GDNQ catalytic center). We conclude that escape from ERDRP-0519 is mediated by secondary structural effects rather than due to primary resistance, which is unusual for non-nucleoside analog polymerase inhibitors (51-53). Conceivably, these substitutions could shift the spatial preference of intrusion and priming loops in the pre-initiation polymerase complex (32), favoring orientation of the intrusion loop towards the cavity wall that is incompatible with compound binding. Naturally, conformational flexibility of the loops cannot be eliminated entirely without loss of polymerase bioactivity. Three observations support the model that resistance arises from a shift in conformational equilibrium of the loops: i) the maximally achievable degree of resistance is moderate and escape can be readily overcome through increased compound concentrations; ii) the loss in compound bioactivity due to resistance (17 to 77-fold increase in EC_50_ depending on the individual mutation) closely resembles the drop in target binding affinity (35 to 63-fold increase in K_D_); and, iii) escape from ERDRP-0519 in all cases carries a substantial viral fitness penalty (10), consistent with disturbed, but not obliterated, polymerase function.

Most notably, resistance mutations greatly eased the ERDRP-0519 block of *de novo* initiation of RNA synthesis in *in vitro* RdRP assays, but had only a marginal effect on restoring RNA elongation in primer extension. This phenotype reveals that restoring RNA synthesis initiation in the presence of compound constitutes the dominant selective force for resistance, identifying inhibition of *de novo* polymerase initiation at the promoter as the primary antiviral effect of ERDRP-0519 in the infected cell. Pharmacological interference with RNA elongation likely represents an artifact of the *in vitro* RdRP assay, due to the use of short and non-encapsidated synthetic template RNA that does not fully recapitulate physiological conditions. The predicted docking pose of ERDRP-0519 in the template RNA channel is consistent with the notion that under physiological conditions the compound cannot engage and arrest a committed polymerase complex in RNA elongation mode. Further experimental support comes from the unchanged steepness of the viral mRNA transcription gradient in infected cells exposed to ERDRP-0519, confirming that the polymerase complex is not susceptible to inhibition any longer once RNA elongation has started.

The ERDRP-0519 docking pose into MeV L provides a direct molecular explanation for the morbillivirus-specificity of the compound, since L microdomains forming the physical binding site of the inhibitor show considerable variability between different genera in the paramyxovirus family (10, 12). To date, native structural information for paramyxovirus polymerases is available only for PIV-5 L, and the resolution of the structure prevents atomic localization of individual side chains (32). Although these restrictions make structure-guided ligand design to broaden the indication spectrum of the ERDRP-0519 chemotype challenging, the insight gained in this study generates high confidence that function-equivalent druggable sites exist in all paramyxovirus, and potentially in all mononegavirus, polymerases. ERDRP-0519 is a clinical anti-MeV candidate that meets fundamental requirements of a valid morbillivirus therapeutic (10, 54). In addition to its direct anti-MeV impact, the compound emerged as a class-defining prototype inhibitor that has illuminated an attractive druggable site in mononegavirus L proteins, revealed a unique inhibitory mechanism of RdRP activity, and has established a foundation for future indication spectrum expansion through scaffold engineering or targeted drug discovery.

## Methods

### Cells and viruses

African green monkey kidney epithelial cells (CCL-81; ATCC) stably expressing human signaling lymphocytic activation molecule (Vero-hSLAM) were maintained at 37°C and 5% CO_2_ in Dulbecco’s modified Eagle’s medium (DMEM) supplemented with 7.5% fetal bovine serum. Insect cells (SF9) were propagated in suspension using Sf-900 II SFM media (Thermo Scientific) at 28°C. All cell lines used in this study are routinely checked for mycoplasma and microbial contamination. All transfections were performed using GeneJuice transfection reagent (Invitrogen), unless otherwise stated.

### Molecular biology

For minigenome studies, resistance mutations were cloned into MeV-Edm L plasmids under T7 control using PCR mutagenesis. All plasmids were sequenced to confirm resistance mutations and sequence integrity. The baculovirus protein expression systems used for L_1708_-P complexes and RdRP assays, were generated as previously described (28).

### Chemical synthesis

All materials were obtained from commercial suppliers and used without purification, unless otherwise noted. Dry organic solvents, packaged under nitrogen in septum sealed bottles, were purchased from EMD Millipore and Sigma-Aldrich Co. Reactions were monitored using EMD silica gel 60 F_254_ TLC plates or using an Agilent 1200 series LCMS system with a diode array detector and an Agilent 6120 quadrupole MS detector. Compound purification was accomplished by liquid chromatography on a Teledyne Isco CombiFlash RF+ flash chromatography system. NMR spectra were recorded on an Agilent NMR spectrometer (400 MHz) at room temperature. Chemical shifts are reported in ppm relative to residual CDCl_3_-*d*_6_ signal. The residual shifts were taken as internal references and reported in parts per million (ppm). To synthesize ERDRP-0519_az_, precursor compound **1** (0.46 gm, 1.0 mmol) (supplementary figure 6) (11) was dissolved in anhydrous DMF (10 ml) in a 50 ml RBF and cooled to 0°C. Under inert atmosphere, 60% NaH in mineral oil (0.06 gm) was added and stirred for 10 minutes. 1-azido-4-(bromomethyl)benzene (0.318 gm, 1.5 mmol) was added and continued stirring at room temperature for 5 hours. After completion, the reaction mixture was quenched with methanol and the solvent was removed under reduced pressure. The crude product was dissolved in dichloromethane (50 ml) and extracted with water (50 ml) and brine (50 ml). The organic layer was dried over anhydrous Na_2_SO_4_, filtered and concentrated under reduced pressure. The crude product was purified by flash column chromatography using ethyl acetate and hexane as eluent. Pure ERDRP-0519_az_ (supplementary figure 6) was obtained as colorless solid, yield 67% (0.4 gm).

^1^H NMR (CDCl_3_, 400 MHz): δ 7.78 (d, *J* = 8 Hz, 2H), 7.19 (d, *J* = 8 Hz, 2H), 7.06 (d, *J* = 8 Hz, 2H), 6.94 (d, *J* = 8 Hz, 2H), 5.79 (s, 1H), 5.04 (s, 2H), 4.21−4.16 (m, 4H), 3.84−3.64 (m, 3H), 3.03−2.96 (m, 1H), 2.76−2.73 (m, 1H), 1.99-1.91 (m, 1H),1.57−1.36 (m, 5H), 1.15−1.07 (m, 1H), 0.97−0.87 (m, 1H). ^19^F NMR (376 MHz, CDCl_3_) δ -62.31; ^13^C NMR (100 MHz, CDCl_3_) δ 159.61, 145.18, 141.56, 140.14 (q, *J* = 38 Hz), 140.05, 136.18, 132.29, 130.28, 130.14, 128.41, 128.28, 120.45 (q, *J* = 268 Hz), 119.45, 119.31, 107.16 (d, *J* = 17 Hz), 58.30, 53.17, 49.54, 49.41, 40.97, 40.15, 40.02, 32.31, 27.88, 24.16, 18.31. MS (ES-API) [M+Na]^+^ : 614.0

### Minigenome dose-response assays

MeV firefly luciferase based minigenomes were performed as described (28). BSR-T7/5 cells (1.1 × 10^4^ per well in a 96-well plate format) were transfected with N-Edm, P-Edm, L-Edm or L-Edm resistant mutant variants, and the MeV luciferase replicon reporter, followed by incubation with three-fold serial dilutions of compound. Luciferase activities were measured using a BioTek Synergy H1 multimode microplate reader approximately 40 hours after transfection. Raw data were normalized according to norm. value [%] = (RLU_sample_ - RLU_min_) / (RLU_max_ - RLU_min_)*100, with RLU_max_ representing vehicle (DMSO) volume equivalent-treated transfected wells and RLU_min_ representing transfected wells lacking the L polymerase-encoding plasmid. Four-parameter variable slope regression modeling was used to determine inhibitory concentrations (EC_50_).

### Antiviral dose-response assays

Compound was added in three-fold dilutions to 96-well plates seeded with Vero-hSLAM cells (1.1 × 10^4^ per well), followed by infection (MOI = 0.2 TCID_50_ units per cell) with a recombinant MeV-NanoPEST reporter virus harboring Nano luciferase (28). After 30-hour incubation, Nano luciferase signals were quantified using a BioTek Synergy H1 multimode plate reader. Raw data were normalized according to norm. value [%] = (RLU_sample_ - RLU_min_) / (RLU_max_ - RLU_min_)*100, with RLU_max_ representing vehicle (DMSO) volume equivalent-treated infected wells and RLU_min_ representing infected wells exposed to 1 mg/ml cycloheximide. Four-parameter variable slope regression modeling was used to determine EC_50_ and EC_90_ concentrations.

### Cytotoxicity testing

To determine cytotoxic concentrations Vero-hSLAM cells were plated in 96 well plates (1.1 × 10^4^ per well) and incubated with three-fold serial dilutions of compound. After incubation of uninfected cells for 72 hours, PrestoBlue substrate (Invitrogen) was added to quantify cell metabolic activity as described (55, 56). Signal was measured using a BioTek H1 synergy multimode microplate reader. Where applicable, cytotoxic concentrations (CC_50_) were calculated based on four-parameter variable slope regression modeling.

### qRT-PCR analyses of viral and cellular transcripts

For MeV transcripts, Vero-hSLAM cells were infected with recMeV-Anc (MOI = 3 TCID_50_ units/cell) by spin-inoculation (2000 x g, 30 min, 4°C) and incubated in the presence of ERDRP-0519 or vehicle at 37°C for 4 hours. Total RNA was extracted four hours after infection using Trizol and subjected to reverse transcription using either oligo-dT primers. Subsequent qPCR used primer pairs specific for MeV P mRNA, MeV H mRNA, MeV L mRNA, or human GAPDH mRNA, respectively. qPCR was performed using an Applied Biosystems 7500 Real-Time PCR System with a StepOnePlus Real-Time PCR System. Samples were normalized for GAPDH.

### MeV polymerase complex expression and purification

MeV P and L proteins were expressed in the Fastbac dual expression system in SF9 cells as previously described (28). Approximately 76 hours after infection, cells were lysed in buffer containing 50 mM NaH_2_PO_4_, 150 mM NaCl, 20 mM imidazole, pH 7.5, 0.5% NP-40 buffer. MeV L_1708_-P and MeV L-P complexes were purified by Ni-NTA affinity chromatography Protein complexes were eluted using buffer containing 50 mM NaH_2_PO_4_, pH 7.5, 150 mM NaCl, 0.5% NP-40, and 250 mM imidazole, and buffer were subsequently exchange to 150 mM NaCl, 20 mM Tris-HCl, pH 7.4, 1 mM DTT and 10% glycerol by dialysis or with Zeba desalting spin columns.

### Biolayer interferometry

MeV L_1708_ constructs were expressed and purified as previously described (28). EZ-Link NHS-Biotin (ThermoFisher) was used to biotinylate MeV L_1708_ samples. Subsequently, buffer was exchanged to Octet Kinetics buffer (Fortebio) using a Zeba desalting spin column. Biotinylated MeV L_1708_ preparations were then bound to Super Streptavidin (SSA) high-binding biosensors (Fortebio) and incubated with increasing concentrations of ERDRP-0519 (0.1 μM to 200 μM) with alternating incubations in Kinetics buffer to generate small-molecule binding and dissociation curves. SSA sensors loaded with mouse anti-FLAG IgG were used as reference sensors and additional biocytin blocked sensors were used as double parallel controls for nonspecific binding. Local fitting of binding kinetics was used to determine K_D_ values using the Octet Red software package by Pall ForteBio. To visualize whether saturation of binding was reached, concentration-dependent steady-state sensor response signals were plotted.

### *In vitro* MeV RdRP assay

For *de novo* polymerase assays, purified P-L hetero-oligomers containing approximately 20 ng of L protein were diluted in 50 µl reaction buffer containing 1 mM DTT, 10% glycerol, 50 mM Tris/Cl (pH 7.4) and mixed with 3 mM MnCl_2_, 1 mM of GTP, UTP and CTP, 50 µM ATP, 10 µCi of α-^32^P labelled ATP (Perkin Elmer), 1 µM of a 16-nt RNA template and 5% DMSO containing ERDRP-0519 at the specified concentration. The 16-nt RNA template sequence 3’-UGGUCUUUUUUGUUUC 5’ (Dharmacon), was derived from the MeV Trailer complement promoter sequence with an additional UUUUU insertion that has been reported to promote polyadenylation for paramyxoviral polymerases (28, 46). Reactions were incubated at 30°C for 5 hours, RNAs ethanol-precipitated overnight at -20°C, resuspended in 50% deionized formamide with 10 mM EDTA, heat-denatured at 95°C for 5 minutes, and fractionated through 20% polyacrylamide gels containing 7 M urea in Tris-borate-EDTA buffer. Polymerization products were visualized through autoradiography with either X-ray films (CL-XPosure, ThermoFisher Scientific) or exposed to BAS Storage Phosphor Screens MS 2040 and scanned on a Typhoon FLA7000 imager (GE Healthcare) for densitometric quantifications. *De novo* polymerase assays were performed on a 25-nt RNA template derived from the RSV trailer complement promoter 3’-UGCUCUUUUUUUCACAGUUUUUGAU 5’ (35) (Dharmacon), using the same transcription buffer as above with the following adjustments: 4 mM MnCl_2_, 2 µM 25-mer RNA template, 1 mM ATP, 50 µM GTP and 10 µCi of α-32P labelled GTP (Perkin Elmer).

### *In vitro* primer extension MeV RdRP assay

For primer extension assays, purified P-L hetero-oligomers containing approximately 100 ng of L protein were diluted in 5 µl reaction buffer containing 1 mM DTT, 5% glycerol, 10 mM Tris/Cl (pH 7.4) and mixed with 3 mM MnCl_2_, 10 µM of either GTP and ATP or GTP only as specified in figure captures, 2.5 µCi of α-32P labelled GTP (Perkin Elmer), 4 µM of 16-nt RNA template, 100 µM of 5’-phosphorylated 4-nt primer ACCA (Perkin Elmer) and 5% DMSO containing ERDRP-0519 at the specified concentration. Reactions were incubated at 30°C for 5 hours, stopped with one volume of deionized formamide with 20 mM EDTA, and analyzed through electrophoresis and autoradiography as described above.

### Photoaffinity labeling-based target mapping

Purified MeV L_1708_ was incubated with 40 µM ERDRP-0519_az_ for 15 minutes prior to activating the crosslinker. The MeV L1708 -ERDRP-0519_az_ mixture was incubated on ice and exposed to UV light (365 nm) for 30 min. The sample was then exposed to additional UV light (254 nm) for 15 minutes. Protein was then collected using FLAG resin. Bound resin was then incubated with Laemmli buffer at 56°C for 15 minutes. SDS-PAGE electrophoreses was then performed using Laemmli buffer on 4-15% acrylamide gels. Bands of interest were excised and analyzed by mass spectrometry. ERDRP-0519_az_ crosslinked peptides were identified by the Proteomics & Metabolomics Facility at the Wistar Institute as previously described (28). In order to identify ERDRP-0519_az_ crosslinked peptides, mass addition of 563.1814 (MW of ERDRP-0519_az_) was considered for all amino acid residues.

### 3D-QSAR model building

The MOE software package (MOE 2018.1001 (57)) was used to perform all conformation searches, energy minimization, and model building. 3D-QSAR modeling was performed using the AutoGPA module embedded in MOE (38), as previously reported (27). For the creation of predictive 3D-QSAR models, a set of 33 diverse analogs reported in (11, 13, 58) were chosen exhibiting various inhibitory activities (EC_50_ concentrations from 0.005-65 µM). This set of compounds was divided into a training set (16 entries;15a-i, 15k, 15m-p, 15r, 16677) and a test set (17 entries; 1a, 28c, 2g, 2i, 2k, 3a, 3-d, 9a, AS-105, AS-136, MS-12, MS-26, MS-27, ERDRP-519az, and ERDRP-0519). Inhibitory concentrations (EC_50_) were converted to pIC50 (−log_10_(EC_50_)). Conformational libraries of each compound were generated using the conformational search function of the AutoGPA package as previously reported (27). A total of 2398 and 3485 conformations were generated for the training and test sets, respectively. Training set conformations were then aligned and assigned pharmacophore features using the pharmacophore elucidation function of MOE. AutoGPA identified common features and created 287 initial pharmacophore models based on the training set. After electrostatic, steric, and space filling model building was performed, a partial-least squares analysis was performed and each model was validated, scored, and ranked based on LOO correlation (q^2^) and goodness of fit (R^2^). The predictive potential of the model was then tested using a conformation database of the test set. Predicted pIC50 values were then compared with actual pIC50 values to determine the correlation (R^2^) and slope of the correlation.

### Homology modeling, *in silico* ligand docking, and pharmacophore extraction

Docking studies were performed with MOE 2018.1001, using the Amber10 force field. Homology models of MeV L based on the coordinates reported for PIV-5 L (PDB 6v85), RSV-L (PDB 6pzk), VSV L (PDB 5a22), and RABV L (PDB 6ueb) were used for docking studies. After protonation and energy minimization, an induced-fit protocol was used to dock the ERDRP-0519 in the MeV L structure derived from PIV-5 L. Target selection was based on resistance data and crosslinking results. Initially, residues H589, S768, T776, 952-966 (crosslinked peptide 1), 1151-1167 (crosslinked peptide 2), L1170, R1233, and V1239 were selected to identify the target site for docking ERDRP-0519, AS-136 (58), and 16677(13). All favorable docking poses were localized proximal to residues 1155-1158. These residues were selected as the target for *in silico* docking and an additional round of binding was performed for ERDRP-0519, AS-136, and 16677. Results from this docking procedure identified a favorable shared docking pose. For covalent docking of ERDRP-0519_az_, residues 1155-1158 were chosen as the reactive site and an induced-fit covalent protocol was used to dock the ERDRP-0519_az_ structure into MeV L.

### Statistical analysis

Excel and GraphPad Prism software packages were used for data analysis. One-way or two-way ANOVA with Dunnett’s, multiple comparisons post-hoc test without further adjustments were used to evaluate statistical significance when more than two groups were compared or datasets contained two independent variables, respectively. The specific statistical test applied to individual studies is specified in figure legends. When calculating antiviral potency and cytotoxicity, effective concentrations were calculated from dose-response data sets through 4-parameter variable slope regression modeling, and values are expressed with 95% confidence intervals (CIs) when available. Biological repeat refers to measurements taken from distinct samples, and results obtained for each biological repeat are shown in the figures along with the exact sample size (n). For all experiments, the statistical significance level alpha was set to <0.05, exact P values are shown in individual graphs.

## Acknowledgements

We thank H-Y Tang and the Wistar Institute Proteomics and Metabolomics Facility for assistance with proteomics analysis and KK Conzelmann for the BSR-T7/5 stable cell line. This work was supported, in part, by Public Health Service grants AI071002 (to RKP) and AI153400 (to RKP), from the NIH/NIAID. The funders had no role in study design, data collection and interpretation, or the decision to submit the work for publication.

## Supplementary Figures

**Supplementary Figure 1**. Location of ERDRP-0519 resistance mutations (black) in comparison with escape sites from an RSV L capping inhibitor (blue) and RSV L blocker AZ-27 (magenta). Side views of the MeV L complex and top view of the RdRP and capping domains only are shown. The GDNQ catalytic center is highlighted in red.

**Supplementary figure 2**. Crosslinking-identified peptide 3 (orange spheres) is located in a highly disordered region of the MeV L polymerase. Localized residues directly engaged by ERDRP-0519_az_ are shown as dark orange spheres. Peptides 1 (pink) and 2 (purple) are also highlighted.

**Supplementary Figure 3**. Synthetic evolution of the ERDRP-0519 chemotype does not alter the pharmacophore. **A-F)** Top scoring *in silico* docking poses (orange sticks) and 2-D ligand interaction projections are shown for the clinical candidate ERDRP-0519 (A-B), first generation lead AS-136 (C-D), and original screening hit 16677 (E-F). Conserved hydrogen bond interactions are predicted between all scaffolds and Y1155, N1285, and H1288. Locations of peptide 2 (purple sticks), of the intrusion (blue) and priming (red) loops, and of proposed PRNTase motifs (blue sticks) are shown. **G)** Stick model overlays of the docking poses of ERDRP-0519 (orange), AS-136 (red) and 16677 (blue).

**Supplementary Figure 4**. *In silico* docking attempts of ERDRP-0519 into PIV-5, RSV and VSV L polymerases, which are all not inhibited by the compound. **A-F)** Docks of ERDRP-0519 into PIV-5 L (red sticks (A-B)), RSV L (red sticks (C-D)) and VSV L (red sticks (E-F)) did not yield poses similar to that obtained for docking into MeV L (orange sticks). 2-D schematics of the top scoring docking poses are shown for each structure. Peptide 1 (pink), peptide 2 (purple), the intrusion (blue) and priming (red) loops, ERDRP-0519 resistance mutations (black spheres), and the proposed PRNTase motifs (blue spheres) are marked.

**Supplementary Figure 5**. Conformation rearrangements of the priming and intrusion loops into initiation conformation eliminates ERDRP-0519 binding pocket. **A-I)** Overlay of the ERDRP-0519 docking pose into MeV L homology models based on PIV-5 (intrusion loop down; A, D, G), RABV (intrusion loop up; B, E, H), and RSV (intrusion loop up; C, F, I) L proteins. Close up views of the ERDRP-0519 binding pocket in the different MeV L models are shown in (D-F). Steric incompatibilities (red scaffold substructures) are noted with RABV and RSV-based models (H-I) with intrusion loop in initiation conformation.

**Supplementary Figure 6**. Chemical synthesis strategy of ERDRP-0519_az_.

